# Tracking changes in functionality and morphology of repopulated microglia in young and old mice

**DOI:** 10.1101/2024.07.12.603244

**Authors:** Zuzanna M. Luczak-Sobotkowska, Patrycja Rosa, Maria Banqueri Lopez, Natalia Ochocka, Anna Kiryk, Anna M. Lenkiewicz, Martin Furhmann, Aleksander Jankowski, Bozena Kaminska

## Abstract

Microglia (MG) are myeloid cells of the central nervous system supporting its homeostasis and instigating neuroinflammation in pathologies. Single-cell RNA sequencing (scRNA-seq) revealed the functional heterogeneity of MG in mice brains. Inhibition of colony-stimulating factor 1 receptor (CSF1R) signaling with inhibitors deplete microglia which rapidly repopulate. Functionalities of repopulated microglia are poorly known. We combined scRNA-seq, bulk RNA-seq, immunofluorescence and confocal imaging to study functionalities and morphology of repopulated microglia. CSRF1R inhibitor (BLZ-945) depleted MG in 21 days and their numbers were restored 7 days later as evidenced by TMEM119 staining and flow cytometry. ScRNA-seq and computational analyses demonstrate that repopulated MG originate from preexisting MG progenitors and reconstitute functional clusters but upregulate inflammatory genes. Percentages of proliferating, immature MG displaying inflammatory gene expression increase in aging mice. Morphometric analysis of MG cell body and branching shows distinct morphology of repopulated MG, particularly in old mouse brains. We demonstrate that with aging some repopulated MG fail to reach the homeostatic phenotype. These differences microglia may contribute to the deterioration of microglia protective functions with age.

## INTRODUCTION

Microglia (MG) are myeloid cells in the central nervous system (CNS) which together with perivascular, choroid plexus, and meningeal macrophages play a crucial role in maintaining CNS homeostasis (Kettenmann *et al*, 2013; Li & Barres, 2018). Microglia are instrumental in maintaining brain homeostasis during development, aging, and disease. Fate mapping, mulitple marker imaging and single-cell transcriptomic studies demonstrated that during disease, aging, or injury, microglia undergo context-dependent transcriptional, morphological, and functional reprogramming acquiring beneficial or deleterious functions (Masuda *et al*, 2020; Mathys *et al*, 2019; Ochocka & Kaminska, 2021; Sousa *et al*, 2018).

Microglia are long-living cells and their survival and proliferation are controlled by Colony-stimulating factor-1 (CSF-1) (Green *et al*, 2020). Microglia depletion protocols have been widely exploited to better understand their roles in the CNS homeostasis. CSF1R inhibitor (e.g. PLX339 that crosses the blood-brain barrier, BBB) delivered for several weeks in the diet depletes MG which rapidly repopulate CNS reaching similar cell densities, morphology, and gene expression profiles as resident microglia (Elmore *et al*, 2014). The brain volume, BBB and neuroinflammation markers were unperturbed by short- (7 days) or long-term (2 months) depletion of MG with PLX3397. The absence of microglia did not affect mouse behavior or reduce performance in standard anxiety, locomotion, learning, and memory tasks (Elmore *et al*, 2015). Fate-mapping approaches and parabiosis experiments excluded the blood origin of repopulated microglia and demomonstrated that all repopulated microglia originate from the few surviving microglia by proliferation (Huang *et al*, 2018). A transient nestin expression in new microglia was detected, but none of the repopulated microglia originated from nestin-positive non-microglial cells (Huang *et al*, 2018). Repopulated microglia had larger cell bodies and less complex extension branching, which resolved over time. LPS (lipopolysaccharide)-induced profiles of inflammatory gene expression in repopulated microglia were similar to control ones. Mice subjected to three rounds of depletion/repopulation (7/7 days) reached a complete repopulation; but failed to replenish microglia after one more round of depletion (Najafi *et al*, 2018).

Advances in single-cell RNA sequencing (scRNA-seq) allow to integrate a comprehensive information about different molecular states, infer functional phenotypes and deduce putative relationships between individual cells. ScRNA-seq studies revealed a considerable heterogeneity of brain myeloid cells (mostly microglia) in naïve mice, including several functional states of microglia (Ochocka *et al*, 2021). We sought to determine if the observed MG heterogeneity is reconstituted after repopulation. For this purpose, we employed scRNA-seq to track transcriptomic changes in individual cells and confocal imaging to determine morphology of repopulated microglia in young and old mice. We demonstrate that after depletion the remaining microglia proliferate, replenish the brain after cessation of the CSF1Ri treatment, and restore the functional heterogeneity. However, the reestablishment of the homeostatic MG phenotype is less effective with brain aging and morphological alterations suggest abnormalities of microglia in older age.

## RESULTS

### Repopulated microglia reconstitute transcriptional heterogeneity but show upregulation of inflammatory genes

Mice were feeded daily with 200 mg/kg BLZ495 (delivered in a peanut butter to ensure that they ate a whole amount) (Fig.1A). The brains of 3-weeks old mice were dissected at day 14 or 21 after BLZ495 and 7 days after the treatment cessation. Myeloid cells were immunmosorted using flow cytometry (fluorescent Cx3cr1+ cells) and we evaluated the kinetics of microglia depletion. Representative flow cytometry graphs show changes in the abundance of microglia under different conditions. At day 21 of the treatment, we noted a complete depletion of microglia, and after 7 days without CSF1Ri percentages of microglia were restored to those in control animals (Fig. 1B-C). In parrallel, brains were collected for immunostaining and microglia were visualized with TMEM119 staining (a microglial marker that shows processes and cell bodies). A few TMEM119-positive cells were detected in the brains of BLZ-945-fed mice, mostly fragments of microglial processes are visible. Repopulated microglia are visible on day 7^th^ after stopping BLZ-495 administration (Fi. 1D).

**Figure 1.**
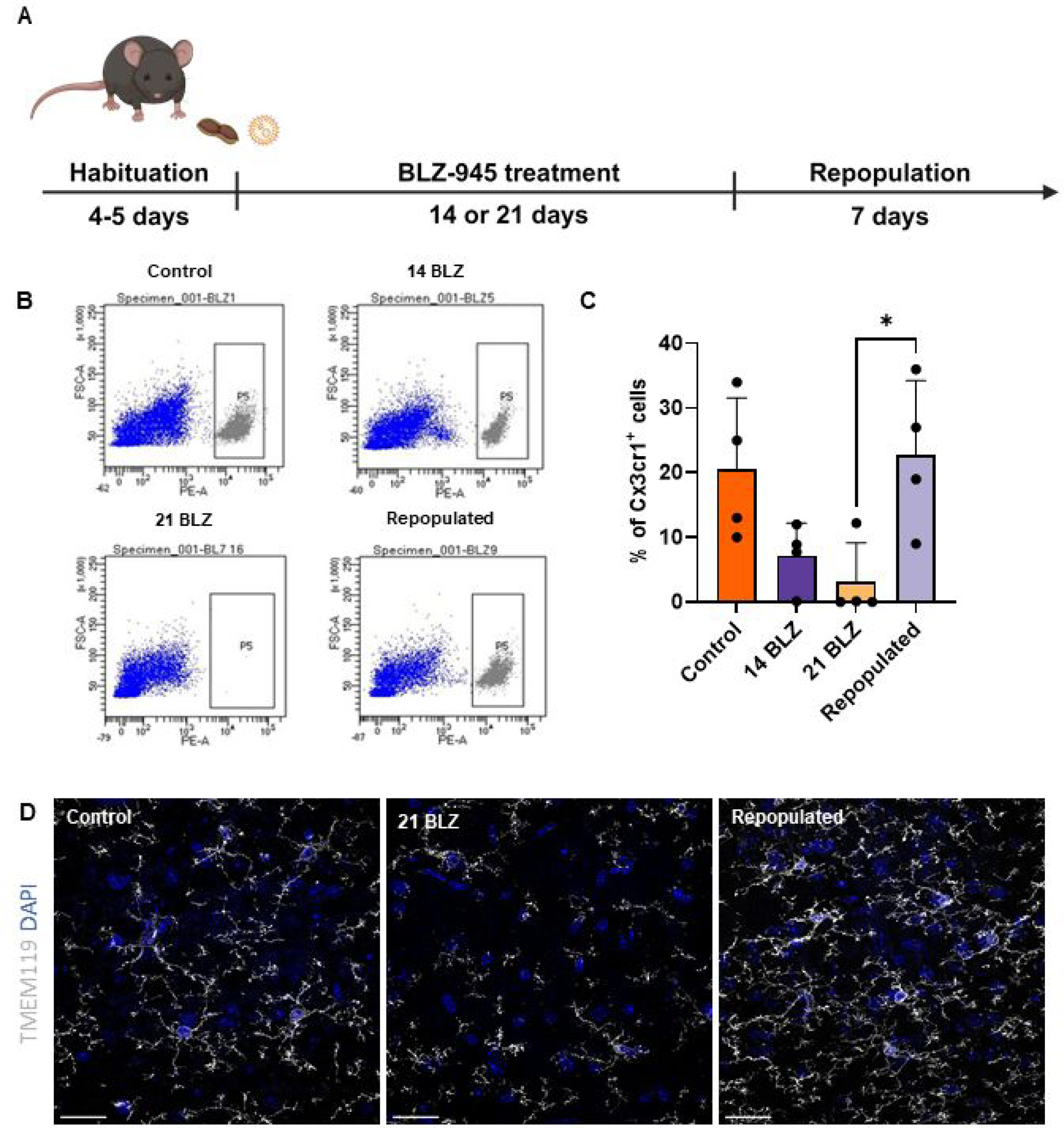
Kinetics of microglia depletion and repopulation. (**A**) Scheme of the experimental workflow. Mice were given orally peanut butter (PB) or BLZ-945 (200 mg/kg) for 14 or 21 days, or were fed with BLZ-945 for 21 days and left for 7 days. (**B**) Representative graphs show quantification of microglia (Cx3cr1+ cells) from the brains of control or BLZ-945 fed mice as determined by flow cytometry. Mice were perfused with PBS, their brains removed and subjected to enzymatic digestion. (**C**) Fluorescent Cx3cr1+ cells were evaluated by flow cytometry (n=4), statistical significance calculated with one-way ANOVA with multiple comparisons (Tukey’s test df =12, p=0.0384). (**D**) TMEM119 staining of microglia in brains of controls, mice fed with BLZ-945 for 21 days, and mice fed with BLZ-945 for 21 days and then left for 7 days. Mice after indicated treatments were perfused with PFA, brains were removed, fixed and subjected to immunofluorescence (IF) staining followed by cell visualization with confocal microscopy. Representative IF images show Tmem119+ microglia in the controls, loss of microglia after 21 days of the treatment, with the remnants of filaments visible, and repopulated, branched microglia 7 days after stopping BLZ-945 administration. Scale bar 50 µm.

ScRNA-seq provides an excellent means to characterize the identities and functionalities of cells under physiological and pathological conditions. We evaluated if the repopulated microglia recreate the functional heterogeneity detected in intact brains and whether these processes undergo similarly in young and aging mice. We performed scRNA-seq on immunosorted CD11b+ cells as previously described (Ochocka *et al*, 2021) from brains in four experimental conditions: control or repopulated in young or aging mice. The cells were filtered and clusters of cells were visualized using Uniform Manifold Approximation and Projection (UMAP) (Fig. 2A). Collectively, we sequenced 30,904 single cells from 16 samples and identified 16 cell clusters/types/states across all conditions. We clustered the cells into 16 cell clusters using Seurat and further merged these clusters (Appendix Fig. S1A) into 8 subtypes based on annotations from previous studies (Ochocka *et al*, 2021, 2023). A majority of immune cells in control and repopulated brains were homeostatic microglia (Fig. 1A). Microglia clusters were investigated in more depth and minor clusters associated with monocytes, macrophages, T cells, NK cells, and Cd24a+ cells were excluded from further analysis (Appendix Fig. S1A).

**Figure 2.**
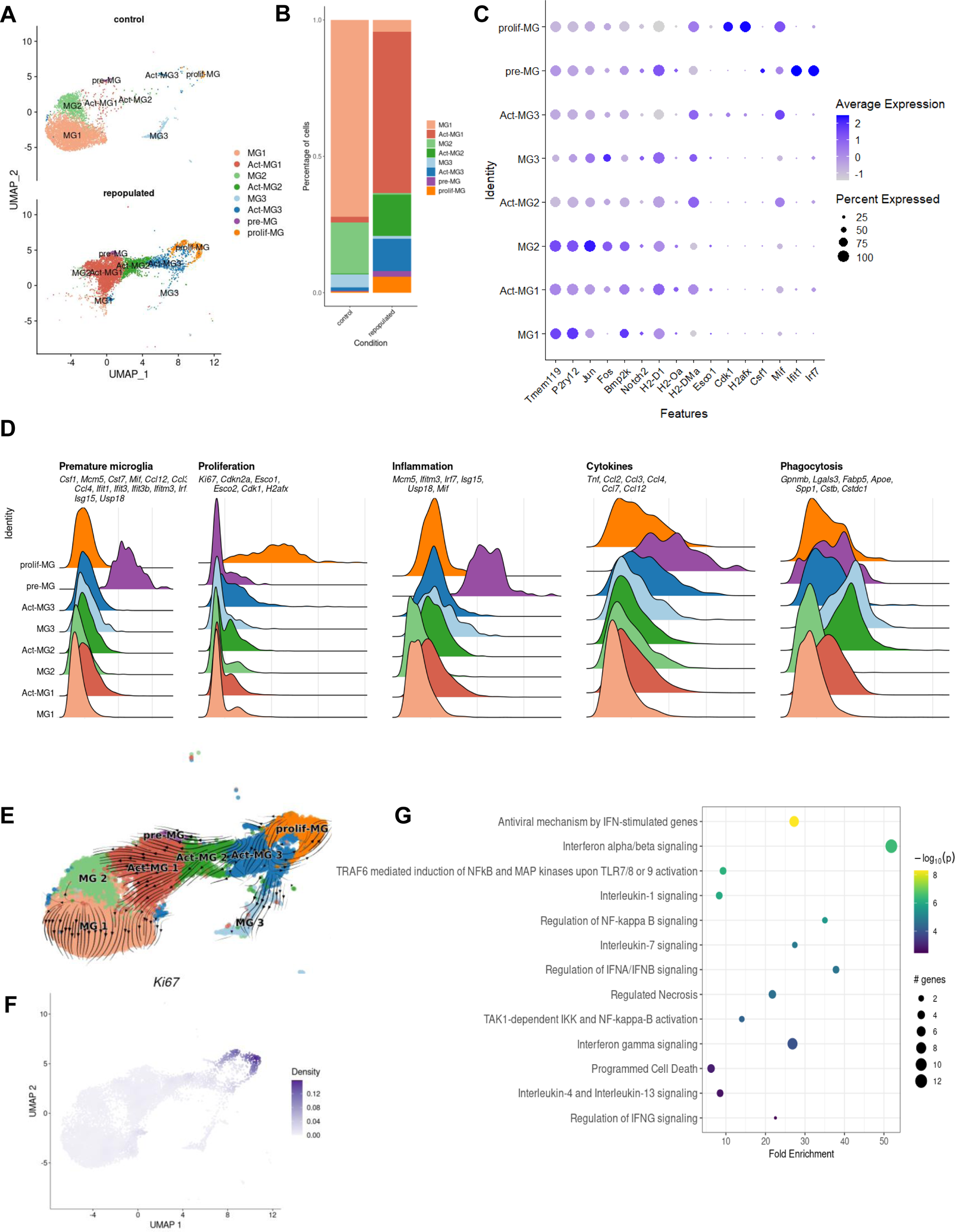
Characterization of the functional heterogeneity of repopulated microglia with single-cell transcriptomics. **(A)** UMAP plot of scRNA-seq data showing cell clustering of control and repopulated microglia. For each condition, four scRNA-seq replicates were combined. Cell clusters are colored and based on the annotations from (Ochocka *et al*, 2021, 2023): MG1 – homeostatic microglia, Act-MG1 – activated homeostatic microglia, MG2 – transcriptionally active cells, Act-MG2 – activated transcriptionally active cells, MG3 – signaling inhibitors and transcriptional repressors, Act-MG3 – activated signaling inhibitors and transcriptional repressors, pre-MG – premature microglia, prolif-MG – proliferating microglia. (**B**) The proportion of cells assigned into each cell cluster in two experimental groups. (**C**) Dot plot showing the average expression levels of signature genes from previous studies (Ochocka *et al*, 2021) for the cell types or processes as defined above. (**D)** Ridge plots showing signature gene score profiles in each cell cluster associated with the specific cell types or processes: premature microglia, proliferation, inflammation, cytokines, and phagocytosis. **(E)** UMAP plot overlaid with RNA velocity vectors (as quantified by scVelo), demonstrating expression dynamic across cell clusters of microglia. Of note, all resulting clusters are derived from proliferating microglia. **(F)** UMAP plot showing weighted kernel density (Alquicira-Hernandez J & Powell J, 2021) of scaled Ki67 gene expression. **(G)** Selected enriched Reactome terms in a set of the differentially expressed genes in the premature microglia (pre-MG) cluster compared to other cell clusters from the Fig. 2A.

The identified clusters MG1, MG2, MG3, Act-MG1, Act-MG2, and Act-MG3 corresponded to Hom-MG1, Hom-MG2, Hom-MG3, Act-MG1, Act-MG2 and Act-MG3 clusters described previously (Ochocka *et al*, 2023). The difference in a subtype composition between repopulated and control brains (Fig. 2B) shows the appearance of repopulated cells in an active state (Act-MG1, Act-MG2 and Act-MG3 clusters), and the emergence of premature and proliferating MG subtypes. The presence of Act-MG1-3 clusters among repopulated microglia is surprising as they resemble functionalities of MG in brains after glioma implantation, and reflect phagocytic, migratory and extracellular matrix degrading activities of activated microglia. The proliferating MG cluster (prolif-MG) is characterized by high expression of the cell cycle genes (*e.g. Ki-67, Esco1/2, Cdk1, H2afx*). Interestingly, the premature MG (pre-MG) cluster shows high expression of inflammation-related (*e.g. Mcm5, Ifitm3, Irf7, Isg15, Usp18, Mif*) and cytokine encoding genes (*Tnf, Ccl2, 3, 4, 7, 12*) suggesting its active role in directing repopulation-related cell migration (Fig. 2A-D).

Trajectory inference has improved scRNA-seq research by enabling the study of dynamic changes in gene expression. It allows the discovery of expression changes that are associated with specific cells in the trajectory, or differentially expressed between distinct differentiation steps (Van den Berge *et al*, 2020). In the reduced dimensional space, a cell pseudotime for a given lineage/state is a distance between a cell state and the origin of the lineage. These advances permit studies of the dynamics of biological processes, such as differentiation patterns from progenitors to more differentiated cellular states (Byrnes *et al*, 2018). Pseudotime ordering identified a trajectory that originates in prolif-MG and through active MG1-3 states leads to homeostatic microglia MG1-3 clusters (Fig. 2E). The prolif-MG cluster shows high expression of *Ki-67* (a marker of proliferation) (Fig.2F). Kyoto Encyclopedia of Genes and Genomes (KEGG) analysis of biological processes in the pre-MG cluster shows high expression of inflammation-related genes of the interferon-, TLR-, NFkB signaling pathways and cytokine encoding genes suggesting their intermediate inflammatory state before they become homeostatic MG (Fig. 2G).

### Defective reconstitution of the homeostatic phenotype by repopulated microglia from older mice

We compared the composition and heterogeneity of repopulated microglia in young (3 months old) and aging (12 months old) mice. The previously identified MG clusters were detected in scRNA-seq samples from young and older mice, and their contribution was similar in naïve mice independently of age (Fig. 3A). However, the contribution of repopulated cells from a specific cluster differed between young and older mice (Fig. 3B). We found increased percentages of active cell states (Act-MG1-3) among repopulated microglia in older mice. The repopulated MG from older mice were enriched in the active MG3 and prolif-MG clusters, while the repopulated MG from young mice had more act-MG1 cells. Trajectory analysis shows that repopulated MG from older mice were enriched in prolif-MG clusters and did not reconstitute intermediate act-MG and homeostatic MG clusters at the same speed as young mice (Fig.3A-C). To compare the expression of selected marker genes across 4 conditions, we plotted the normalized expression levels and a fraction of cells expressing them, separately for each of the 8 cell clusters. We verified that the selected genes had similar expression profiles across the experimental conditions, confirming their suitability as marker genes (appendix S2). The KEGG analysis of genes upregulated in prolif-MG clusters of repopulated MG revealed that the enriched genes are linked to proliferation (mRNA splicing, DNA replication, and chromatin remodeling) (Fig.3D). The profiles of selected differentially expressed genes are shown on the heatmap (Fig.3E).

**Figure 3.**
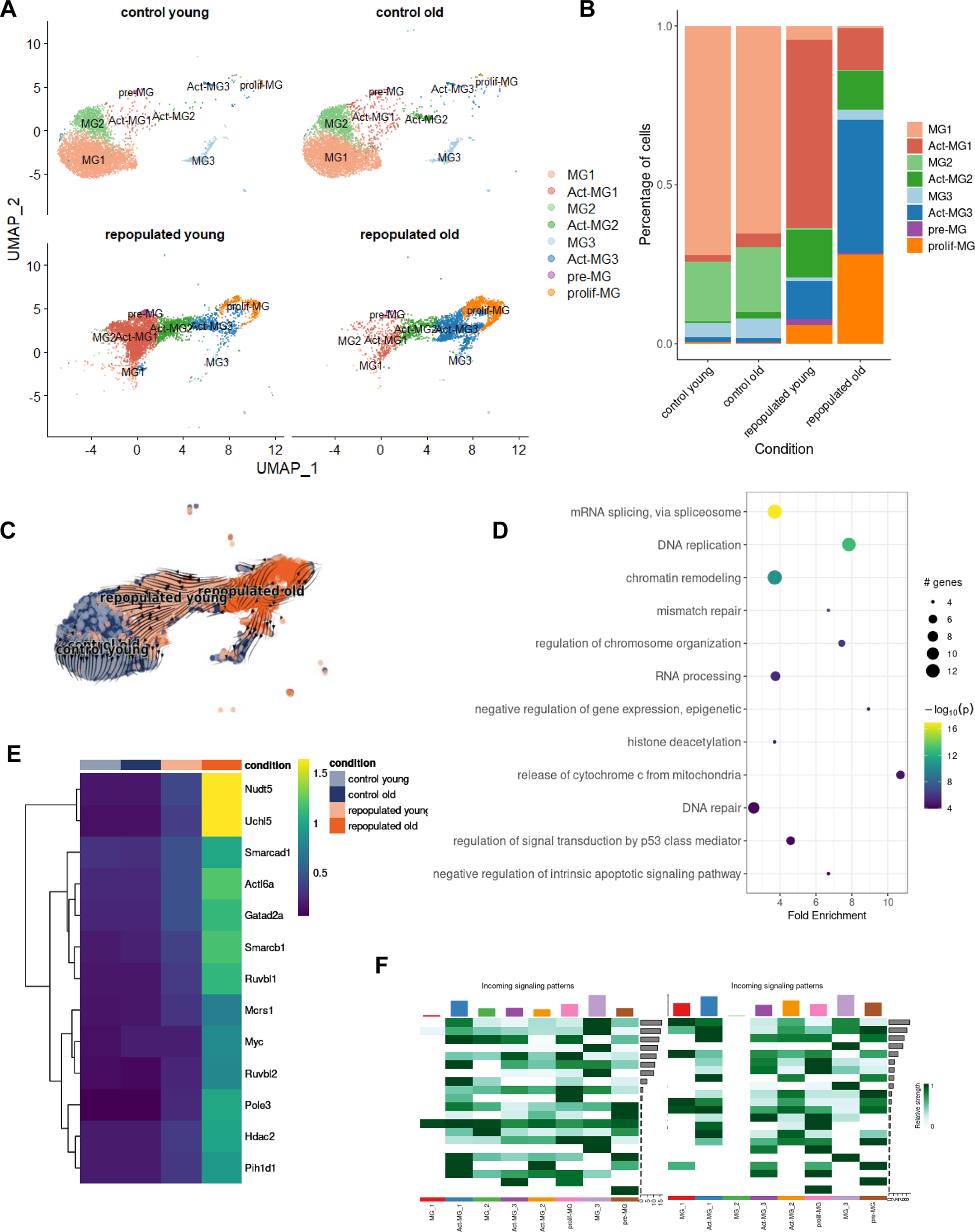
Dysfunctional microglial repopulation in aging mice. (**A**) UMAP plot of scRNA-seq data showing cell clustering obtained for control and repopulated microglia in young and aging mice. For each condition, four scRNA-seq replicates were combined. Cluster annotations as in Fig. 3A. (**B**) Fraction of cells assigned into each cell cluster, separately for each of four conditions. (**C**) UMAP plot overlaid with RNA velocity vectors (as quantified by scVelo), demonstrating expression dynamic across experimental conditions. (**D)** Selected enriched KEGG pathways in the set of differentially expressed genes in the Activated MG3 (Act-MG3) cluster compared to other cell clusters. (**E**) Heatmap of average expression of upregulated genes from the chromatin remodeling KEGG pathway and enriched in Activated-MG3 cell cluster, divided by experimental conditions. (**F)** Communication probability via selected ligand-receptor pairs (y-axis) for each directed pair of cell clusters (x-axis).

Identification of ligand-receptor pairs in scRNA-seq data can be used to infer intercellular communication from the coordinated expression of their cognate genes (Efremova *et al*, 2020). We explored the dataset searching for expression of the genes associated with the interacting proteins in the gene expression matrix and used those values as inputs to compute a communication score for each ligand-receptor pair using a scoring function of the CellPhoneDB v.2.0. Putative R–L interactions between various subpopulations of repopulated microglia show abundant outgoing interactions from pre-MG and act-MG1 involving genes coding for galectins, CCLs, ICAM, semaphorin 4, MIF. C-C motif chemokine ligands (CCLs) are known drive the chemotaxis of myeloid and lymphoid cells (Gschwandtner *et al*, 2019). Galectins (Soares *et al*, 2021) and intercellular cell adhesion molecule (ICAM)-1 (Lyck & Enzmann, 2015) control the migration of immune cells in brain parenchyma during inflammation. Macrophage migration inhibitory factor (MIF) is a pleiotropic protein, participating in inflammatory and immune responses (Nishihira, 2000). Similar subpopulations have increased incoming signaling patterns (Fig.3F).

The observed differences in the dynamics of repopulated microglia in aging mice prompted us to examine transcriptional patterns of control and repopulated microglia from brains of aged mice (18-22-month-old). We analyzed bulk RNAseq data from CD11b+ immunosorted cells and performed the comparison between repopulated and control groups using linear regression models. The sample variance was estimated using Principal Component Analysis dimension reduction and one sample, which was a significant outlier compared to the other samples, was excluded. The volcano plot shows (Fig. 4A) a number of differentially expressed genes (DEGs) in microglia from old mice. A KEGG analysis of genes upregulated in repopulated microglia in old mice unravels some pathways, which are enriched among upregulated DEGs. Interestingly, several of those pathways are related to cell death, apoptosis, senescence, HIF1, Ca^2+^, FoxO signaling (Fig.4B). The heatmap illustrates the expression of the top 50 DEGs which shows that numerous genes expressed in homeostatic microglia in controls have lower expression in repopulated microglia (Fig.4C). We evaluated the expression of selected genes from those pathways in scRNAseq dataset and found that expression of proapoptotic genes (*Lmnb1, Casp8, Psmd3, Bak1*) is increased in repopulated MG from older mice (Fig.3D). The expression of genes related to the apoptosome and cell death markers is restricted to the acti-MG3 cluster in older mice (Fig.3E). We explored other gene signatures upregulated in microglia during aging, neurodegenerative, or neuroinflammatory processes, and found that many of these genes are upregulated in repopulated MG, in particular from older mice. The observed differences in transcriptomic profiles of repopulated microglia suggest the emergence of the transient inflammatory phenotype, particularly in microglia from old mice. With age, repopulated microglia display elevated expression of apoptosis-related genes which is indicative that some cells may undergo cell death.

**Figure 4.**
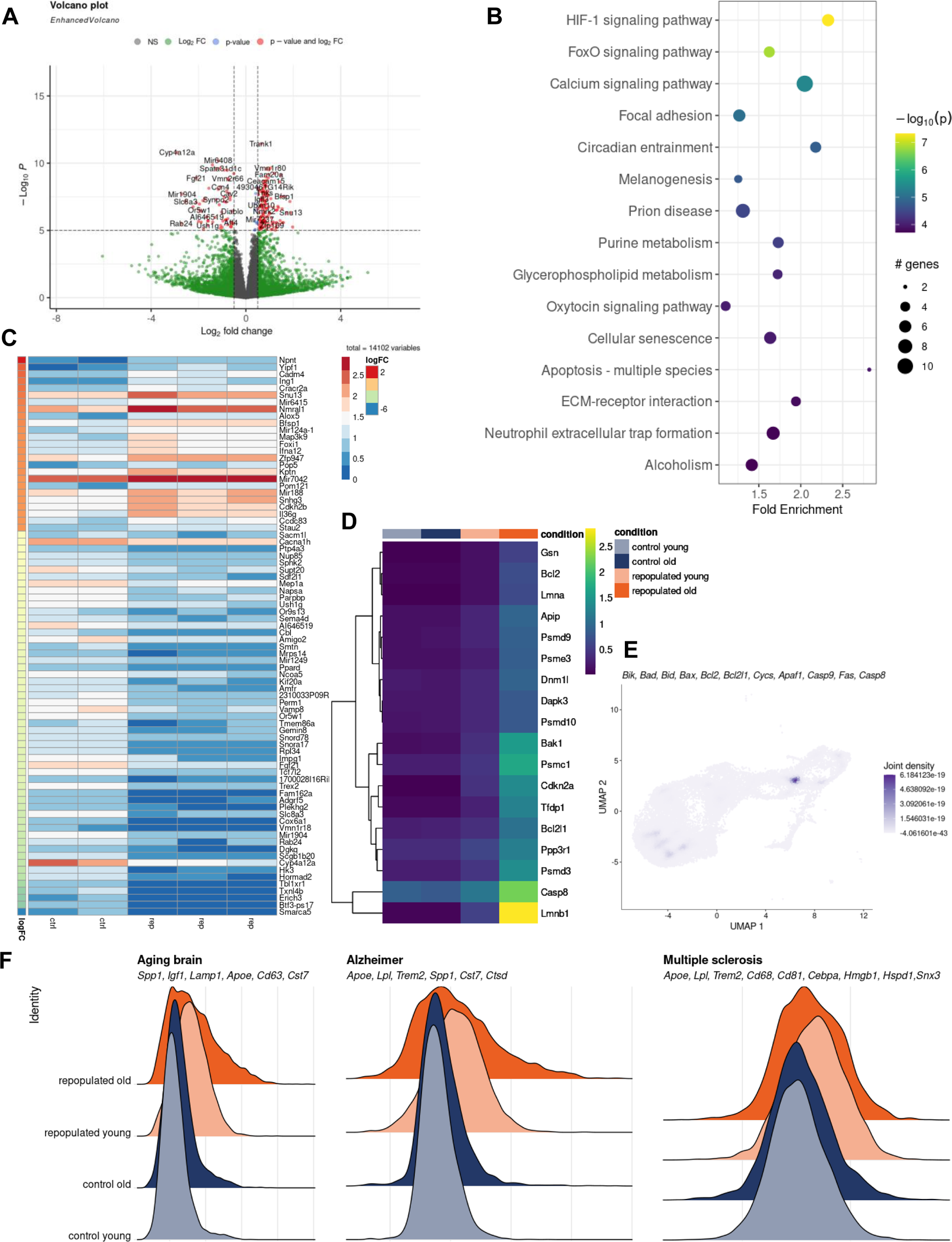
Transcriptional characteristics of repopulated microglia from aged mice. Volcano plot showing expression fold change for all genes in control and repopulated microglia from aged mice (18-22 months old), along with *p*-value for differential expression. Mice were given orally PB or BLZ-945 (200 mg/kg) for 21 days and left for 7 days. Mice were perfused with PBS and their brains were removed, subjected to enzymatic digestion, and CD11b+ cells immunosorted by FACS, total RNAs were isolated and subjected to RNAseq (n=4/group). (**B**) Top 15 enriched KEGG pathways in the set of differentially expressed genes (p-value and log_2_ FC criteria) from the Fig. 4A. (**C**) Heatmap of top 50 differentially upregulated genes with the highest logFC value and adjusted p-value < 0.05 in the bulk RNAseq dataset. (**D**) Heatmap of average expression of the upregulated genes from “Apoptosis – multiple species” KEGG pathway in scRNA-seq data. (**E**) UMAP plot showing weighted kernel density of scaled gene expression of selected apoptosis marker genes in sc-RNAseq data from the fig. 3A. (**F**) Ridge plots showing expression scores of selected microglial neurodegeneration genes characteristics for the aging brain, Alzheimer’s disease and multiple sclerosis (from Paolicelli *et al*, 2022) in the scRNA-seq dataset.

### Repopulated microglia in older mice display different morphology than control microglia

Microglia in the CNS show mostly ramified morphology typical for mature cells with homeostatic functions (Tay *et al*, 2017). Even unchallenged microglia show frequent extension and motility of microglial processes (Davalos et al, 2005; Nimmerjahn *et al*, 2005). Confocal microscopy and 3D imaging with specialized programs such as IMARIS Software allow tracing of cell morphology and quantification of various parameters such as cell volume, length of microglial processes, a number of terminal points, branch points, and number of cell segments (Bosch & Kierdorf, 2022). The changes in gene expression reported above suggested different functikonality of repopulated microglia in the aging brain. We performed immunofluorescence staining for TMEM119 and evaluated morphology of repopulated, TMEM119+ microglia in the brains of young (2-3 months) and old (16-18 months) mice using confocal microscopy and 3-D reconstruction of cell shapes. Repopulated microglia show distinct and more complex morphology than control ones, with more numerous branches and more elongated microglial processes with more branch points (Fig.5A). Using Scholl analysis, the radius of MG branches was quantified and we found more cells with higher radius among repopulated microglia in brains of young mice. Such differences were not observed among repopulated microglia in brains of old mice that showed less complex branching (Fig.5B, C). We determined more morphological parameters and found that repopulated microglia in young mice have a bigger area but branching and filament lengths are similar to microglia in control mice. In contrast, repopulated microglia in old mice have significantly shorter filaments and less branching (Fig.5D). These results show that the morphological complexity of microglia is restored after repopulation in the brains of young but not old mice.

**Figure 5.**
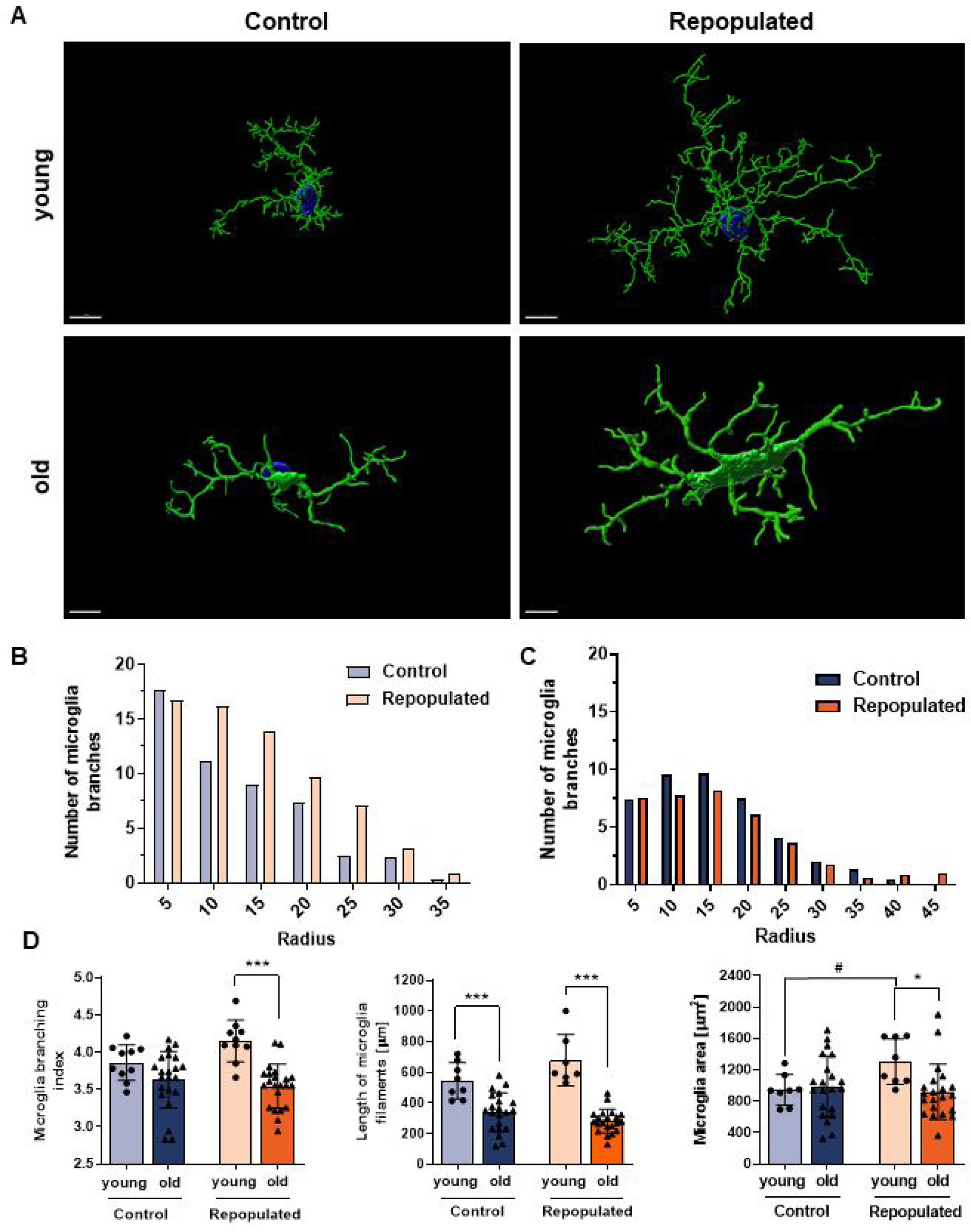
Different morphology of repopulated microglia from aged mice. (**A**) Representative confocal microscopy images of Trem1919+ microglia. 3D models created with the Imaris Software (Oxford Instruments, UK). Scale bar 7 µm. Of note, in young mice repopulated microglia are more branched, with numerous elongated filaments with secondary processes located distally from the cell body. Microglia after repopulation have bigger cell bodies. In old mice (18-22 months) repopulated microglia are branched similarly to controls and have less branches/processes than in the brains of young mice. (**B, C**) Quantification of control or repopulated microglia branching with a Sholl analysis (radius 5 µm) in young (**B**) and old (**C**) mice, respectively. (**D**) Comparison of the microglia branching (branching index and length of filaments) in young and old mice (for branching, Sidak’s Test df = 60, p=0.0001, for the length of microglial filaments, Sidak’s Test control group df=53, p=0,0001, repopulated group df=53, p=0,0001); the area covered by Trem119+ microglia (Sidak’s Test df=54, p=0.0193).

## DISCUSSION

Microglia states are defined by their intrinsic and extrinsic determinants, as well as their spatiotemporal context. The intrinsic determinants (ontogeny, sex or genetic background) as well as age, spatial location, and environmental factors impact MG states (Paolicelli *et al*, 2022). While core markers define microglia identity as a subpopulation of myeloid cells distinct from other brain myeloid cells and peripheral macrophges, they are not informative about the functional “state” of microglia, which depends on the context. MG are exceptionally responsive to alterations in their local environment, and their morphology, ultrastructure and molecular profile are dynamic, which results in many different cell states that could be resolved with single-cell technologies (Ochocka *et al*, 2021; Paolicelli *et al*, 2022;). Herein, we present the scRNAseq-based characterization of MG states after repopulation induced by pharmacological depletion of MG with the CSF1R inhibitor. We demonstrate that repopulated MG recapitulates all states/phenotypes occurring in control MG, which is shown by the presence of the all identified scRNAseq clusters. However, we found the manifestation of transiently activated microglia phenotypes (Act-MG1-3) that resemble proliferating, phagocytic, migratory microglia detected in brain tumor niches (Ochocka *et al*, 2023). This is not unexpected, as newly arising MG must migrate through the parenchyma to various regions of the brain, degrade extracellular matrix on the way, and this migration is driven by cytokines and chemokines. Newly generated microglia, arising from precursor MG (pre-MG), display elevated expression of inflammation-related genes (mostly from interferon-related pathways) that may reflect mild inflammation associated with repopulation before mature microglia resume their homeostatic functions. The ligand-receptor interaction analysis shows the putative signals outgoing from pre-MG to other clusters, which highlights its decisive role in driving repopulation. We confirmed that all new MG clusters originate from the remaining pre-MG.

Under control conditions, the same MG states are detected in the brains of young and aging mice. However, among the repopulated cells from the brains of aging mice we detecet more proliferating MG that do not reach mature states as MG in the brains of young animals. The results sugegst that those microglia do not reach the homeostatic states with the same dynamics as in brains of young mice. Comparison of bulk RNA-seq results on repopulated CD11b+ cells from brains of young and aged mice (16-18 months old) shows pronounced downregulation of homeostatic MG genes and upregulation of genes related to cell death and apoptosis in microglia from aged mice. Exploration of scRNAseq results shows that while repopulated MG from young mice augment expression of inflammation-related genes (which is abrogated in the homeostatic MG1-3 clusters), in repopulated MG from older mice gene signatures upregulated in microglia during aging, neurodegenerative or neuroinflammatory processes are elevated and those cells do not mature into homeostatic microglia (reduced MG1-2 clusters). The results point to the deficiency of MG from older and aged mice to achieve homeostatic phenotypes, which might impact their functions in supporting neuronal homeostasis. Age has a key influence on the microglial homeostatic state, as age-dependent changes in microglia are characterized by an enrichment of defined regulatory factors and gene expression profiles (Hammond *et al*, 2019; Li *et al*, 2019). MG can adopt an aberrant dystrophic or senescent phenotype, and aged microglia have been frequently identified in neurodegenerative diseases (Antignano *et al*, 2023; Angelova & Brown, 2019). Aged MG upregulates MHC, CD86, CD68, CR3 proteins and pattern recognition receptors such as TLRs and Clec7a (Niraula *et al*, 2017).

Microglial morpohology is a good indicator of their states due to morphological transformation of activated microglia. We demonstrate that while there is much difference in morphology of microglia in young and aged mice, except for shorter processes in the latter, after repopulation microglia from aged mice show shorter processes and less branching indicative of alterations of their functions. Pronounced morphological changes of microglia such as deramification, cytoplasmic fragmentation, and shortening of cellular processes in the aged brain (even in the absence of forthright pathology) have been detected in human postmortem brain samples (Streit & Xue, 2010; Streit *et al*, 2004). As microglia are both immunological defenders and neuroprotective cells in the CNS, the consequences of microglia dysfunction might involve increased susceptibility to brain infections and a failure to cope with neurodegenerative processes.

## MATERIALS AND METHODS

### Animals and treatment

All experimental procedures on animals were approved by the First Local Ethics Committee for Animal Experimentation in Warsaw (#836/2019; #1364P1/2022). Young (3 months old), aging (12 months old) or aged (18-22 months old) female and male GFP-M::Cx3cr1-CreERT2fl/fl::Rosa26-tdTomatotg/tg mice were housed in the Nencki Institute, Poland. Animals were kept in individually ventilated cages, with free access to food and water, at the temperature of 21–23 °C, 50–60% humidity, under a 12 h/12 h day and night cycle. Animals were habituated for 3-4 days by being placed individually in a cage for 3 h, where a small amount (∼100 µL) of the peanut butter (PB) was provided. Mice were fed with BLZ-945 (MedChemExpress, HY-12768) at a dose of 200 mg/kg b.w. daily in 100 µL PB (Peanut butter, GoOn, Santé); controls received PB. BLZ-945 was delivered for 14 or 21 days to establish the kinetics of depletion; repopulation was studied 7 days after cessation of the treatment.

### Tissue dissociation and isolation of CD11b+ by flow cytometry

Mice were anesthetized with 3% isoflurane (Iso-Vet), overdose of ketamine (Biowet Pulawy), and xylazine (Sedazin, Biowet) (160 mg/kg and 20 mg/kg of body weight, respectively) injected i.p., followed by perfusion with cold PBS. The brains were collected and placed in cold HBSS (without Ca^2+^ and Mg^2+^). In the first experiment, a whole hemisphere was isolated for flow cytometry experiments and the remaining hemisphere was stored in cold PFA for 48 h. Subsequently, brains were transferred into 30 % sucrose (w/v) in PBS for 48 h and frozen in Tissue Freezing Medium (Leica) at −80°C.

For flow cytometry, tissue was minced, transferred onto c-tubes containing 1,950 µL DMEM, and supplemented with 50 µL of Deoxyribonuclease I (Sigma Aldrich). Dissociation was performed with the MACS dissociator with heaters (Militenyi Biotec) for 22 minutes and later the samples were passed through 70 nm and 40 nm strainers into 50 mL cold HBSS with ions to stop the enzymatic reaction (EasyStrainer™, BioOne). The cell suspension was centrifuged (10 min, 4°C, 300xg), the supernatant was removed and the pellet was resuspended in 25 ml of Myelin Gradient Buffer with 5 ml of cold PBS overlayed, followed by centrifugation (950xg, 4°C, 20’ with acc. 0 and deacc. 0). The pellet was re-suspended in 1 mL of BD buffer (Dulbecco’s Phosphate-Buffered, Stain Buffer, FBS, BD Pharmigen™). Cells were counted using the Nucleocounter (Chemometec). For flow cytometry, cells were suspended in the Stain Buffer (BD Pharmingen) with anti-mouse CD16/CD32 Fc Block™ 1:200 (BD Pharmigen) for 10 min. Next, anti-mouse CD11b antibody 1:250 (M1/70, BD Pharmigen) was added and cells were incubated for 20 min at 4 °C, then washed with Stain Buffer. Cells were sorted using purity precision mode on FACS Aria (BD FACSAria Cell sorter BD-Biosciences) into 200 μL of sterile PBS. The sorted sample was mixed by inverting the Eppendorf tubes and placed on ice, for no longer than 30 minutes.

### Total RNA isolation from CD11b+ cells

For RNA isolation, sorted CD11b+ cells were stored in the −80°C freezer. The samples were thawed on ice and total RNA was isolated with the RNAeasy Plus Mini kit (Qiagen) according to the manual. RNA concentration was immediately determined using a spectrophotometer NanoDrop2000 (ThermoScientific).

### Single-cell RNA-seq (scRNA-seq) experiments

Directly after sorting, cell quantity and viability of CD11b^+^ cells were determined and a cell suspension volume containing 5,000 target cells was used for further processing. Preparation of gel beads in emulsion and libraries were performed with Chromium Controller and Single-Cell Gene Expression v2 Chemistry (or v3 for naïve vs sham-implanted experiment – Supplementary Figure 1) (10x Genomics) according to the Chromium Single-Cell 3’ Reagent Kits v2 (or v3) User Guide provided by the manufacturer. Libraries’ quality and quantity were verified with a High-Sensitivity DNA Kit (Agilent Technologies, USA) on a 2100 Bioanalyzer (Agilent Technologies, USA). Next, sequencing was run in the rapid run flow cell and paired-end sequenced (read 1 – 26 bp, read 2 – 100 bp) on a HiSeq 1500 (Illumina, San Diego, CA 92122 USA). To minimize batch effects, four scRNA-seq libraries (Rep1 to Rep4) were constructed, each combining 4 animals across 4 experimental conditions (control young, control old, repopulated young, repopulated old). For each scRNA-seq library, samples from individual experimental conditions were tagged using TotalSeq™ anti-mouse hashtag antibodies. The hashtag assignment was as follows: control young – #Ab 1, control old – #Ab 2, repopulated young – #Ab 3, repopulated old – #Ab 4:

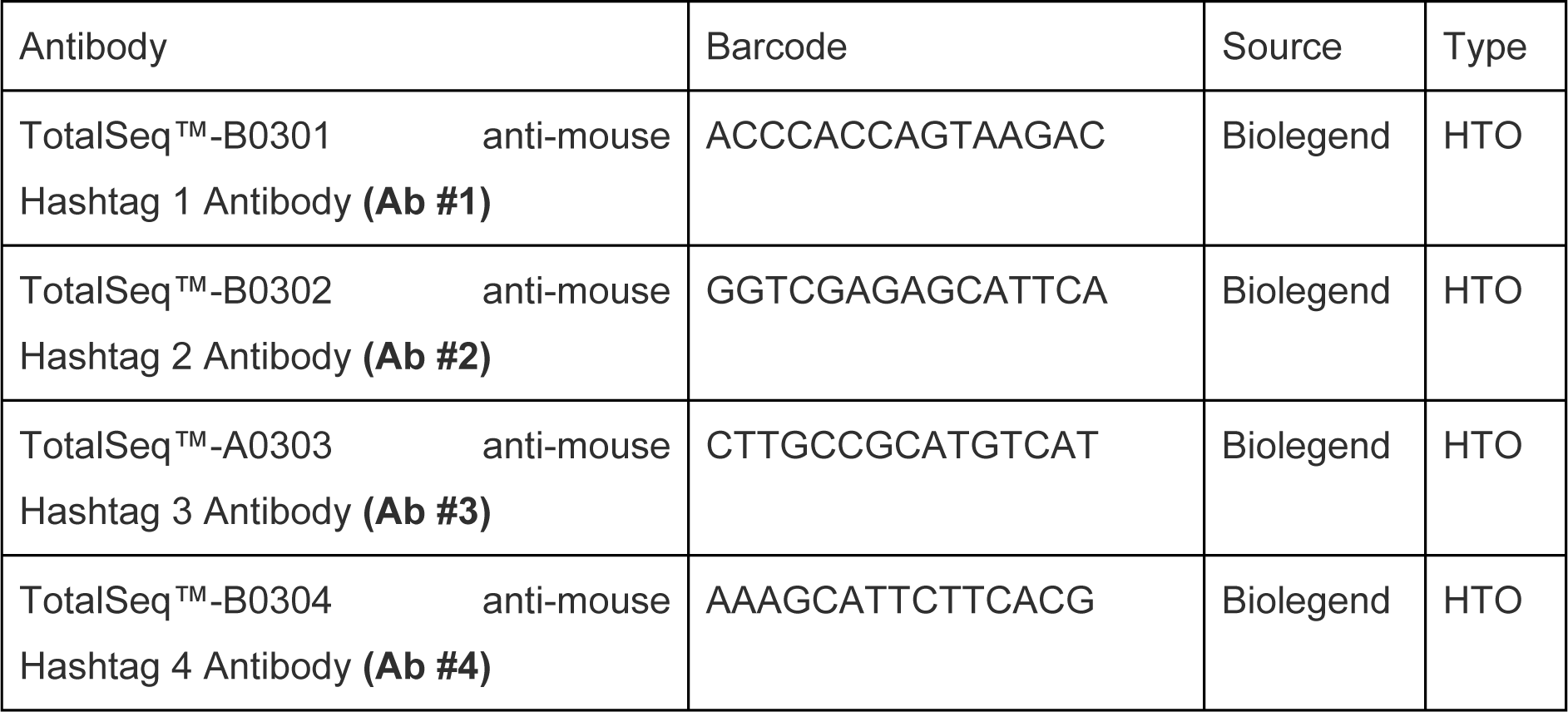

### RNA sequencing

In single-cell and bulk RNAseq experiments, quality and integrity of total RNA were assessed with an Agilent 2100 Bioanalyzer using an RNA 6000 Nano Kit (Agilent Technologies, Ltd.). For bulk RNA-seq strand-specific polyA-enriched RNA libraries were prepared using the KAPA Stranded mRNA Sample Preparation Kit according to the manufacturer’s protocol (Kapa Biosystems, MA, USA). Briefly, mRNA molecules were enriched from 500 ng of total RNA using poly-T oligo-attached magnetic beads (Kapa Biosystems, MA, USA). Obtained mRNA was fragmented and the first-strand cDNA was synthesized using a reverse transcriptase. Second cDNA synthesis was performed to generate double-stranded cDNA (dsDNA). Adenosines were added to the 3′ ends of dsDNA and adapters were ligated (adapters from NEB, Ipswich, MA, USA). Following the adapter ligation, the uracil in a loop structure of the adapter was digested by the USER enzyme from NEB (Ipswich, MA, USA). Adapters containing DNA fragments were amplified by PCR using NEB starters (Ipswich MA, USA). Library evaluation was done with Agilent 2100 Bioanalyzer using the Agilent DNA High Sensitivity chip (Agilent Technologies, Ltd.). The mean library size was 300 bp. Libraries were quantified using a Quantus fluorometer and QuantiFluor double-stranded DNA System (Promega, Madison, Wisconsin, USA). Libraries were run in the rapid run flow cell and were paired-end sequenced 2×76 bp on HiSeq 1500 (Illumina, San Diego, USA) in a case of scRNAseq and 2×151 bp on NovaSeq 6000 (Illumina, San Diego, USA) in a case of RNAseq.

### Single-cell RNA-seq data preprocessing and normalization

Raw scRNA-seq sequencing data (1 671 million raw reads, BCL files) were demultiplexed into libraries (Rep1 to Rep4) and converted to FASTQ files using CellRanger v3.0.1 (10x Genomics) (https://support.10xgenomics.com/single-cell-gene-expression/software/pipelines/latest/installation) and bcl2fastq v2.20.0.422 (Illumina). Sequencing results were mapped to the mouse genome GRCm38 (mm10) downloaded from the 10x Genomics website and quantified using CellRanger. A total number of cells identified by the CellRanger was 30,904. The median number of detected genes per cell was 2,310, and the median unique molecular identifiers per cell was 68,228.

Further data analysis was performed in R using Seurat v4.2.0. (Hao *et al*, 2021) The data was first demultiplexed by their hashtag oligonucleotides, which specified the experimental condition. According to the hashtag oligonucleotides, duplicates (two cells in one droplet) and negatives (empty droplets) were filtered out; the thresholds of finding duplicates and negatives were set for each hashtag separately using MULTIseqDemux() function. The libraries were first processed independently, following a standard Seurat workflow. Unless otherwise specified, all other quantitative parameters were set to default values.

To filter out possible empty droplets and low-quality cells, all cells with <200 transcripts were excluded from the analysis. In addition, dead cells or cells of poor quality, recognized as the ones with >5% of their transcripts coming from mitochondrial genes, were excluded from the downstream analysis. After applying these filters, 19,997 cells were present in the data set. Gene expression measurements for each cell were normalized by the total number of transcripts in the cell using the “LogNormalize” method of the NormalizeData function multiplied by a default scale factor of 10,000, and the normalized values were log-transformed. For each library, the 2,000 most highly variable genes were identified, using variance stabilizing transformation (“vst”). Finally, the libraries were merged into a single Seurat object using the merge() function.

The normalized counts were scaled using Seurat function *ScaleData()*, with all genes given as “features” argument. The dimensionality reduction was performed in the Seurat package with the PCA algorithm in the first 30 dimensions and data visualization in the 2D space was performed with the UMAP algorithm. Clustering was obtained using a method, which constructs a shared nearest neighbor graph by calculating the neighborhood overlap between every cell and its k.param nearest neighbors by using FindNeighbours(). FindClusters() was used with 0.5 resolution, which produced 16 separated cell clusters, by identifying clusters of cells by a shared nearest neighbor (SNN) modularity optimization-based clustering algorithm (Louvain method).

### Single-cell RNA-seq downstream analysis

For follow-up downstream analysis, we identified cell populations present in the data. The comparison was made using ProjecTILs-3.0 R library (Andreatta *et al*, 2021) on the dataset from (Ochocka *et al*, 2023). To further compare the experimental conditions, we set differentially expressed genes using FindMarkers() with negative binomial regression (test.use=”negbinom”) in each condition, comparing corresponding cell clusters. These genes were used to look for the enriched Gene Ontology terms, pathways from KEGG and Reactome databases to identify processes, which were activated or suppressed in compared populations. Enrichment analysis and visualizations were made using pathfindR (v.2.3.1) (Ulgen *et al*, 2019) library, which uses protein-protein interaction network to find significant paths, which activates during ongoing processes. To obtain an interaction network from the dataset, genes with their p-value are mapped into a subnetwork space and sorted according to the score they gain and a number of significant genes. The sorted list of significant genes was used for an enrichment analysis. Enriched terms with adjusted p values larger than the given threshold were discarded and those with the lowest adjusted p value for each term were kept. The process was computed multiple times in parallel to obtain stable results. Enriched terms had computed the distances between themselves and then hierarchical clustering was implemented.

### Trajectory interference

To investigate the dynamics in the microglia repopulation processes, and identify trajectories of cell differentiation, we used scVelo (v0.3.2) library (Bergen *et al*, 2020). We compared the amount of mRNA and kinetics by using ratio between spliced and unspliced fragments. First, velocyto (v.1.0) was used to calculate spliced and unspliced count matrices. Additional meta-data was exported from the Seurat analyzed object. The velocity of RNA was estimated using dynamical modeling. The obtained results were mapped on the UMAP plot.

### Identification of significant ligand-receptor pairs

To interfere with knowledge about the differences between conditions, we used CellChat R library (v.1.5.0) (Jin *et al*, 2023) to analyze intercellular communication from microglia repopulation process data using a combined social network analysis and pattern recognition, on reduced dimensions of the analyzed dataset. This approach allowed us to show a prediction of incoming and outgoing pattern signals from identified cell clusters, and provide precisely their functionality.

### Bulk RNA-seq analysis

Quality control assessment was performed by FastQC, and the report was generated using MultiQC. Trimming adapters was performed. The sequences were aligned to the mm10/GRCm38 reference genome using STAR built-in functions from pipeline RNA-seq-STAR-deseq2 v2.0.0. (https://github.com/snakemake-workflows/rna-seq-star-deseq2). Preprocessing, such as filtering out genes with null variation, normalization, scaling pseudocounts to counts per million, and dimension reduction to linear visualization was performed in the R environment using edgeR library. To expand information about genes, as different database annotations, gene descriptions were done using biomaRt library. During preprocessing, one sample from the controls was excluded because of significantly higher variance, as an outlier from other samples from the same condition. The linear regression model was fitted to the data using limma library. Comparison was made between repopulated and control groups. After testing the FDR correction of p-value was made. The differentially expressed genes were chosen, if their log2FC absolute value was higher than one, and adjusted p-value was lower >0.05. We have obtained 736 differentially expressed genes, 453 upregulated and 283 downregulated, which were used as an input to the pathfindR function to search for activated or deactivated pathways in the analyzed dataset. We used the KEGG pathway database, Reactome, as well as Gene Ontologies, where we focused on Biological Processes. We selected genes of interests, which expression was shown on boxplots.

### Immunofluorescence

For immunostainings, mice were anesthetized and transcardially perfused using PBS and 4% paraformaldehyde (PFA). The brains were removed, fixed in 4% PFA overnight, and then placed in 30% sucrose for 2-3 days before embedding in Tissue-Tek O.C.T Compound. 10 µm thick sections were cut on a cryostat (Microm HM525, Thermo Scientific) and 3 sections were placed on a glass slide (Polysine™ slides, ThermoScientific). Slides were stored at −80°C for further processing. Cryosections were first blocked in PBS containing 10% donkey serum in a 0.1% Triton X-100 solution for 2 hours and then incubated in a 3% donkey serum buffer overnight at 4 °C with guinea pig anti-Tmem119 antibody (1:1,000, Synaptic System). After that, the sections were washed in PBS and incubated with secondary antibody Alexa Fluor 488 (1:1,000, Life technologies) for 2 hours at room temperature. Finally, sections were mounted in a mounting medium with DAPI (Vectashield Vibrance, Vector laboratories). Images were obtained using a Zeiss LSM800 Airyscan confocal microscope.

To quantify microglia morphology in immunofluorescent images, confocal images (21-μm z-stack at 3-μm intervals, Zeiss 800, 40×/1.3 oil objective) were acquired. The number of cell somas per frame was used to normalize all process endpoints and process length. Sholl analysis was automatically performed using the Imaris Software for each cell by counting the number of intersections between microglia branches and each increasing circle. We determined a number of intersections per cell and maximum branch length. From these parameters, a ramification index was calculated to quantify cell branching density.

### Statistical analysis

For animal and transcriptomic experiments a number of animals was n=4. All quantitative data were analyzed using a one-way ANOVA with a significance level of *p* < 0.05 and were performed using GraphPad Prism (version 9.3).

## Acknowledgements

We would like to thank Bartłomiej Gielniewski from the Sequencing Core Facility. This work was supported by the National Centre for Research and Development, Poland ERA-NET-NEURON/18/2018 (B.K.), JPco-fuND2 Programme no. 2022/04/Y/NZ5/00122, the National Science Centre, Poland (B.K.) the Polish National Agency for Academic Exchange (Polish Returns 2019 (A.J.) and European Research Council ERC-CoG 865618 (M.F.).

## Author Contributions

**Zuzanna Łuczak-Sobotkowska:** Formal analysis; Investigation; Visualization; Methodology; Writing—original draft; **Patrycja Rosa:** Data curation; Formal analysis; Visualization; Methodology; Writing—original draft. **Maria Banqueri Lopez**: Investigation; Methodology; **Natalia Ochocka**: Investigation; Methodology; **Anna Kiryk**: Investigation; Methodology; **Anna M. Lenkiewicz**: Investigation; Methodology; **Martin Furhmann**: Investigation; Methodology; **Aleksander Jankowski:** Data curation; Formal analysis; Visualization; Supervision; Funding acquisition; Methodology; Writing—original draft. **Bozena Kaminska**: Conceptualization; Formal analysis; Supervision; Visualization; Funding acquisition; Methodology; Writing—original draft; Writing—review and editing.

## DATA AVAILABILITY

The single-cell and bulk RNAseq RNA-seq datasets produced in this study are available in the European Nucleotide Archive database (https://www.ncbi.nlm.nih.gov/geo/query/acc.cgi?acc=GSE271559

https://www.ncbi.nlm.nih.gov/geo/query/acc.cgi?acc=GSE271560). The computer code produced in this study is available at https://github.com/rosapatrycja/microglia_repopulation_2024.

## REFERENCES

Alquicira-Hernandez J & Powell J (2021) Nebulosa recovers single-cell gene expression signals by kernel density estimation Bioinformatics 37: 2485–2487

Andreatta M, Corria-Osorio J, Müller S, Cubas R, Coukos G & Carmona SJ (2021) Interpretation of T cell states from single-cell transcriptomics data using reference atlases. Nat Commun 12: 2965

Angelova DM & Brown DR (2019) Microglia and the aging brain: are senescent microglia the key to neurodegeneration? J Neurochem 151: 676–668

Antignano I, Liu Y, Offermann N & Capasso M (2023) Aging microglia. Cellular and Molecular Life Sciences 80: 126

Van den Berge K, Roux de Bézieux H, Street K, Saelens W, Cannoodt R, Saeys Y, Dudoit S & Clement L (2020) Trajectory-based differential expression analysis for single-cell sequencing data. Nat Commun 11: 1201

Bergen V, Lange M, Peidli S, Wolf FA & Theis FJ (2020) Generalizing RNA velocity to transient cell states through dynamical modeling. Nat Biotechnol 38: 1408–1414

Bosch LFP & Kierdorf K (2022) The Shape of μ—How Morphological Analyses Shape the Study of Microglia. Front Cell Neurosci 16: 942462

Byrnes LE, Wong DM, Subramaniam M, Meyer NP, Gilchrist CL, Knox SM, Tward AD, Ye CJ & Sneddon JB (2018) Lineage dynamics of murine pancreatic development at single-cell resolution. Nat Commun 9: 3922

Davalos D, Grutzendler J, Yang G, Kim JV, Zuo Y, Jung S, Littman DR, Dustin ML & Gan WB (2005) ATP mediates rapid microglial response to local brain injury in vivo. Nat Neurosci 8, 752–758

Efremova M, Vento-Tormo M, Teichmann SA & Vento-Tormo R (2020) CellPhoneDB: inferring cell–cell communication from combined expression of multi-subunit ligand–receptor complexes. Nat Protoc 15: 1484–1506

Elmore MRP, Lee RJ, West BL & Green KN (2015) Characterizing newly repopulated microglia in the adult mouse: Impacts on animal behavior, cell morphology, and neuroinflammation. PLoS One 10(4): e0122912

Elmore MRP, Najafi AR, Koike MA, Dagher NN, Spangenberg EE, Rice RA, Kitazawa M, Matusow B, Nguyen H, West BL, et al (2014) Colony-stimulating factor 1 receptor signaling is necessary for microglia viability, unmasking a microglia progenitor cell in the adult brain. Neuron 82: 380–397

Green KN, Crapser JD & Hohsfield LA (2020) To Kill a Microglia: A Case for CSF1R Inhibitors. Trends Immunol 41: 771–784

Gschwandtner M, Derler R & Midwood KS (2019) More Than Just Attractive: How CCL2 Influences Myeloid Cell Behavior Beyond Chemotaxis. Front Immunol 10:2759

Hammond TR, Dufort C, Dissing-Olesen L, Giera S, Young A, Wysoker A, Walker AJ, Gergits F, Segel M, Nemesh J, et al (2019) Single-Cell RNA Sequencing of Microglia throughout the Mouse Lifespan and in the Injured Brain Reveals Complex Cell-State Changes. Immunity 50: 253–271.e6

Hao Y, Hao S, Andersen-Nissen E, Mauck WM, Zheng S, Butler A, Lee MJ, Wilk AJ, Darby C, Zager M, et al (2021) Integrated analysis of multimodal single-cell data. Cell 184: 3573–3587.e29

Huang Y, Xu Z, Xiong S, Sun F, Qin G, Hu G, Wang J, Zhao L, Liang YX, Wu T, et al (2018) Repopulated microglia are solely derived from the proliferation of residual microglia after acute depletion. Nat Neurosci 21: 530–540

Jin S, Plikus M V & Nie Q (2023) CellChat for systematic analysis of cell-cell communication from single-cell and spatially resolved transcriptomics. bioRxiv 10.1101/2023.11.05.565674 [PREPRINT]

Kettenmann H, Kirchhoff F & Verkhratsky A (2013) Microglia: New Roles for the Synaptic Stripper. Neuron 77: 10–18

Li Q & Barres BA (2018) Microglia and macrophages in brain homeostasis and disease. Nat Rev Immunol 18: 225–242

Li Q, Cheng Z, Zhou L, Darmanis S, Neff NF, Okamoto J, Gulati G, Bennett ML, Sun LO, Clarke LE, et al (2019) Developmental Heterogeneity of Microglia and Brain Myeloid Cells Revealed by Deep Single-Cell RNA Sequencing. Neuron 101: 207–223.e10

Lyck R & Enzmann G (2015) The physiological roles of ICAM-1 and ICAM-2 in neutrophil migration into tissues. Curr Opin Hematol 22: 53–59

Masuda T, Sankowski R, Staszewski O & Prinz M (2020) Microglia Heterogeneity in the Single-Cell Era. Cell Rep 30: 1271–1281

Mathys H, Davila-Velderrain J, Peng Z, Gao F, Mohammadi S, Young JZ, Menon M, He L, Abdurrob F, Jiang X, et al (2019) Single-cell transcriptomic analysis of Alzheimer’s disease. Nature 570: 332–337

Najafi AR, Crapser J, Jiang S, Ng W, Mortazavi A, West BL & Green KN (2018) A limited capacity for microglial repopulation in the adult brain. Glia 66: 2385–2396

Nimmerjahn A, Kirchhoff F, Helmchen F (2005) Resting Microglial Cells Are Highly Dynamic Surveillants of Brain Parenchyma in Vivo Science 308: 1314–1318

Niraula A, Sheridan JF & Godbout JP (2017) Microglia Priming with Aging and Stress. Neuropsychopharmacology 42: 318–333

Nishihira J (2000) Macrophage Migration Inhibitory Factor (MIF): Its Essential Role in the Immune System and Cell Growth J Interferon Cytokine Res. 20(9): 751–62

Ochocka N & Kaminska B (2021) Microglia diversity in healthy and diseased brain: Insights from single-cell omics. Int J Mol Sci 22: 1–26

Ochocka N, Segit P, Walentynowicz KA, Wojnicki K, Cyranowski S, Swatler J, Mieczkowski J & Kaminska B (2021) Single-cell RNA sequencing reveals functional heterogeneity of glioma-associated brain macrophages. Nat Commun 12:1151

Ochocka N, Segit P, Wojnicki K, Cyranowski S, Swatler J, Jacek K, Grajkowska W & Kaminska B (2023) Specialized functions and sexual dimorphism explain the functional diversity of the myeloid populations during glioma progression. Cell Rep 42: 1119

Paolicelli RC, Sierra A, Stevens B, Tremblay ME, Aguzzi A, Ajami B, Amit I, Audinat E, Bechmann I, Bennett M, et al (2022) Microglia states and nomenclature: A field at its crossroads. Neuron 110: 3458–3483

Soares LC, Al-Dalahmah O, Hillis J, Young CC, Asbed I, Sakaguchi M, O’neill E & Szele FG (2021) Novel galectin-3 roles in neurogenesis, inflammation and neurological diseases. Cells 10: 3047

Sousa C, Golebiewska A, Poovathingal SK, Kaoma T, Pires-Afonso Y, Martina S, Coowar D, Azuaje F, Skupin A, Balling R, et al (2018) Single-cell transcriptomics reveals distinct inflammation-induced microglia signatures. EMBO Rep 19: e46171

Streit WJ, Sammons NW, Kuhns AJ & Sparks DL (2004) Dystrophic Microglia in the Aging Human Brain. Glia 45: 208–212

Streit WJ & Xue Q-S (2010) The Brain’s Aging Immune System Aging Dis. 1(3): 254–261

Tay TL, Savage JC, Hui CW, Bisht K & Tremblay MÈ (2017) Microglia across the lifespan: from origin to function in brain development, plasticity and cognition. Journal of Physiology 595: 1929–1945

Ulgen E, Ozisik O & Sezerman OU (2019) PathfindR: An R package for comprehensive identification of enriched pathways in omics data through active subnetworks. Front Genet 10: 858

